# The impact of gestational diabetes on functional capacity of the infant gut microbiome is modest and transient

**DOI:** 10.1101/2023.09.14.557457

**Authors:** Ryan V. Chieu, Katharine Hamilton, Paul M. Ryan, Julia Copeland, Pauline W. Wang, Ravi Retnakaran, David S. Guttman, John Parkinson, Jill Hamilton

## Abstract

Gestational diabetes mellitus (GDM) is a metabolic complication that manifests as hyperglycemia during the later stages of pregnancy. In high resource settings, careful management of GDM limits risk to the pregnancy, and hyperglycemia typically resolves after birth. At the same time, previous studies have revealed that the gut microbiome of infants born to mothers who experienced GDM exhibit reduced diversity and reduction in the abundance of several key taxa, including *Lactobacillus*. What is not known is what the functional consequences of these changes might be. In this case control study, we applied 16S rRNA sequence surveys and metatranscriptomics to profile the gut microbiome of 30 twelve-month old infants to examine the impact of GDM during pregnancy. Relative to mode of delivery and sex of the infant, maternal GDM status had only a limited impact on the structure and function of the developing microbiome. While GDM samples were associated with a decrease in alpha diversity, we observed no effect on beta diversity and no differentially abundant taxa. Further, while mode of delivery and sex of infant affected the expression of multiple bacterial pathways, most of the impact of GDM status on the function of the infant microbiome appears to be lost by twelve months of age. These data serve to reassure parents and paediatricians that, at least in high resource settings, while mode of delivery appears to impact function and diversity for longer than anticipated, GDM may not have persistent effects on the function nor composition of the infant gut microbiome.

## INTRODUCTION

The human gut microbiome is increasingly viewed as a key determinant of health, with evidence supporting links to an ever-increasing number of diseases from diabetes and obesity to depression^1,2^. While the microbiome can exhibit dramatic changes over the course of an individual’s life, how it develops over the first three years has a critical impact on determining its future contributions to health and disease^3,4^. Among the most impactful factors that contribute to this development are mode of delivery, use of antenatal or postpartum antibiotics, and diet. Initial colonization is largely driven by mode of delivery, with vaginal deliveries associated with the dominance of *Bifidobacterium, Lactobacillus* and *Bacteroides*^5^, species that experience reduced abundance in infants delivered through Caesarean-section (C-section). Diet also comprises several important developmental milestones in the gut microbiome. As an infant’s diet changes from primarily breast or formula milk feeding with the introduction of solids, the microbiome increases in complexity^6^. While most studies have focused on dynamics of community composition using 16S rDNA sequence surveys, our knowledge of functional changes associated with the microbiome at this critical stage of development is limited.

In addition to these impactful postnatal factors, the metabolic status of the mother during pregnancy has also been found to contribute to the formation of the infant microbiome^7^ and future health status^8^. Gestational diabetes mellitus (GDM) is a relatively common metabolic derangement of pregnancy, occurring in 3-20% of pregnancies^9^ in which mothers exhibit hyperglycemia during the later stages of pregnancy^10^. While hyperglycemia often resolves after birth, it can increase the risk of type 2 diabetes and other metabolic disorders^11^(p),^12^. 16S rDNA gene surveys have shown that the gut microbiota of infants born to mothers with GDM exhibit a significant decrease in alpha diversity^13,14^, together with a loss in abundance of several taxa, including *Lactobacillus* and *Flavonifractor*^15^. While such dysbioses have been postulated as contributory to the increased risk of cardiometabolic diseases, little is known concerning the functional implications of such microbiome shifts.

Whole microbiome RNA sequencing, or metatranscriptomics, is a method of surveying the function of a community of microbes. Previous metatranscriptomics studies focusing on the developing infant gut microbiome have identified characteristics of gene expression of major taxa in the infant microbiome^16–18^. Observations regarding function and expression help to form a more complete picture of the early microbiome and how important factors may shape it. In this study, we used metatranscriptomics to functionally profile the microbial communities associated with stool samples from the 12-month-old infant gut. In addition to examining how the function of these communities respond to factors such as mode of delivery, breast feeding status and sex of the infant, we also examined how exposure to GDM impacts both microbial community structure and function. In consideration of the lack of tools specifically built for statistical analysis of metatranscriptomics data, we also examined how well several different differential expression methods applied to our data^19^. To our knowledge, this is the first study to perform community functional profiling on the infant gut microbiome in the context of GDM.

## RESULTS

### Description of cohort

Women in the late 2^nd^ to early 3^rd^ trimester of their pregnancies were recruited into the study during routine visits to Mount Sinai Hospital, Toronto, Canada. Exclusion criteria includes mothers that are currently diagnosed with diabetes, as this would preclude a diagnosis of GDM. Women were recruited both before and after the 1-hour 50g glucose challenge test (GCT). The cohort was enriched for mothers that failed the GCT, who are more likely to be diagnosed with GDM than those with a passing GCT result. Study participants were thereafter stratified by GDM status as measured through 2 hour, 75g oral glucose tolerance test (OGTT) performed according to criteria defined by the National Diabetes Data Group^9,20^. Subsequent to birth, exclusion criteria for infants were: born either < 37 or > 42 weeks of gestation, twins, having conditions that require frequent or prolonged hospitalization, or having conditions that alter cardiometabolic risk. Stool samples were collected and sent for 16S rDNA profiling (3 months and 1 year) and metatranscriptomics profiling (1 year). Additional clinical measurements included: sex (Female and Male – 11 and 19, respectively), exclusive breastfeeding status (Exclusive and Partial – 8 and 22 participants, respectively), mode of delivery (Caesarean section and vaginal – 10 and 20, respectively), maternal BMI (z-score = 0, 1, 2 or NA – 20,11,7 and 6 participants, respectively) and infant BMI at birth (z-score = −2, −1, 0, 1 or NA – 2, 8, 23, 7 and 1, respectively). Participant characteristics are summarized in Table 1 and Supplemental Table 1.

**Table 1.** Study Demographic Table.

|  |  | GDM | NGT | t-test p-value |
| --- | --- | --- | --- | --- |
|  | GDM Condition | 14 | 16 |  |
| Average | Maternal BMI Pre-Pregnancy | 25.67 | 25.14 | 8.09E-01 |
|  | Weight Gain at the time of the OGT (kgs) | 11.01 | 13.98 | 7.39E-02 |
|  | AUC Glucose | 17.29 | 13.79 | 1.47E-09 |
|  | Weeks of Gestation | 38.63 | 39.37 | 2.06E-02 |
|  | Infant Birth Weight (kgs) | 3.40 | 3.45 | 7.07E-01 |
| Number | *Mode of Delivery* |  |  |  |
|  | Vaginal Delivery | 10 | 10 (9) |  |
|  | Cesarean-section Delivery | 4 | 6 (5) |  |
|  | *Sex* |  |  |  |
|  | Female | 5 | 6 |  |
|  | Male | 9 | 10 (8) |  |
|  | *Exclusive Breastfeeding – 3 months\** |  |  |  |
|  | Exclusive | 5 | 6 |  |
|  | Partial | 9 | 7 |  |
|  | *Exclusive Breastfeeding – 12 months* |  |  |  |
|  | Exclusive | 3 | 5 |  |
|  | Partial | 11 | 11 |  |

\* One 3-month sample did not record breastfeeding status
Bracketed number indicates number in 3-month infants

### GDM has limited impact on the microbial community in the developing infant gut

To examine the impact of GDM on the developing infant gut community, we performed 16S rDNA amplicon sequencing to profile stool samples collected at 3-month and 12-month timepoints after birth. From the 6,124,664 sequenced reads (median 83,769) we identified 512 bacterial OTUs across all 58 samples. Alpha diversity analyses (Shannon Index) revealed an increase in alpha diversity for the 12-month samples relative to the 3-month samples (*P* < 0.001, ANOVA) and that vaginal delivery decreases community diversity at 3 months postpartum (*P* < 0.001, ANOVA) and increases at 12 months (*P* < 0.001, ANOVA; Figure 2b), but not when examining all samples regardless of infant age. Similarly, normal glucose tolerance (NGT) infants (P = 0.038; Figure 2a) and exclusive breastfeeding status (EBF; P < 0.001) were associated with an increase in alpha diversity across all infant ages and at 3 months, but not at 12 months. No other associations between alpha diversity and patient metadata were identified (sex, introduction of solids, BMI or insulin sensitivity).

**Figure 1.**
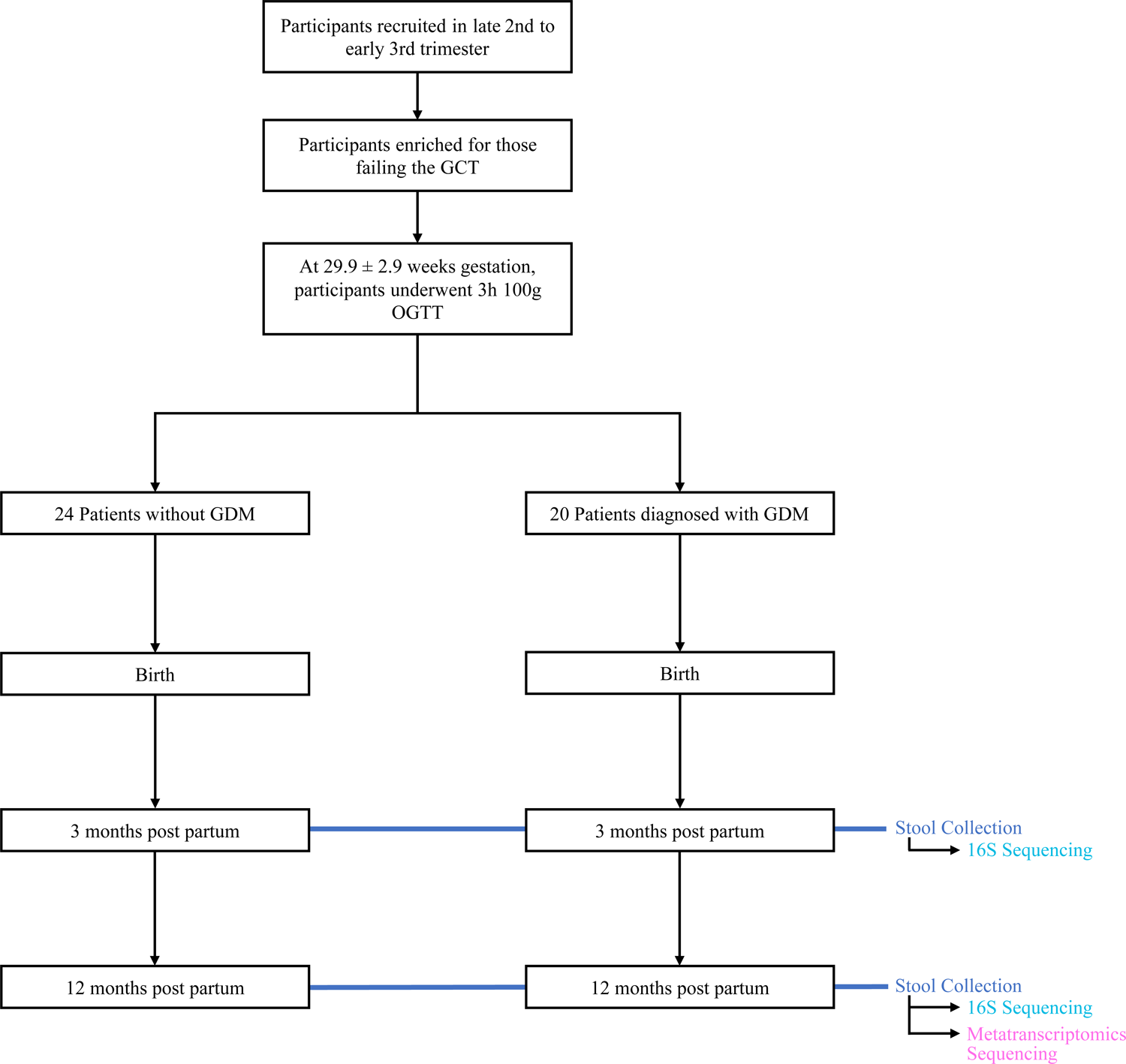
Overview of clinical study design. Patient info was collected at birth, 3 months of infant age, and 12 months of infant age. 16S sequencing was performed on stool samples collected at 3 months and 12 months. Metatranscriptomics sequencing was performed on stool samples collected at 12 months. See Table 1 for demographic information.

**Figure 2.**
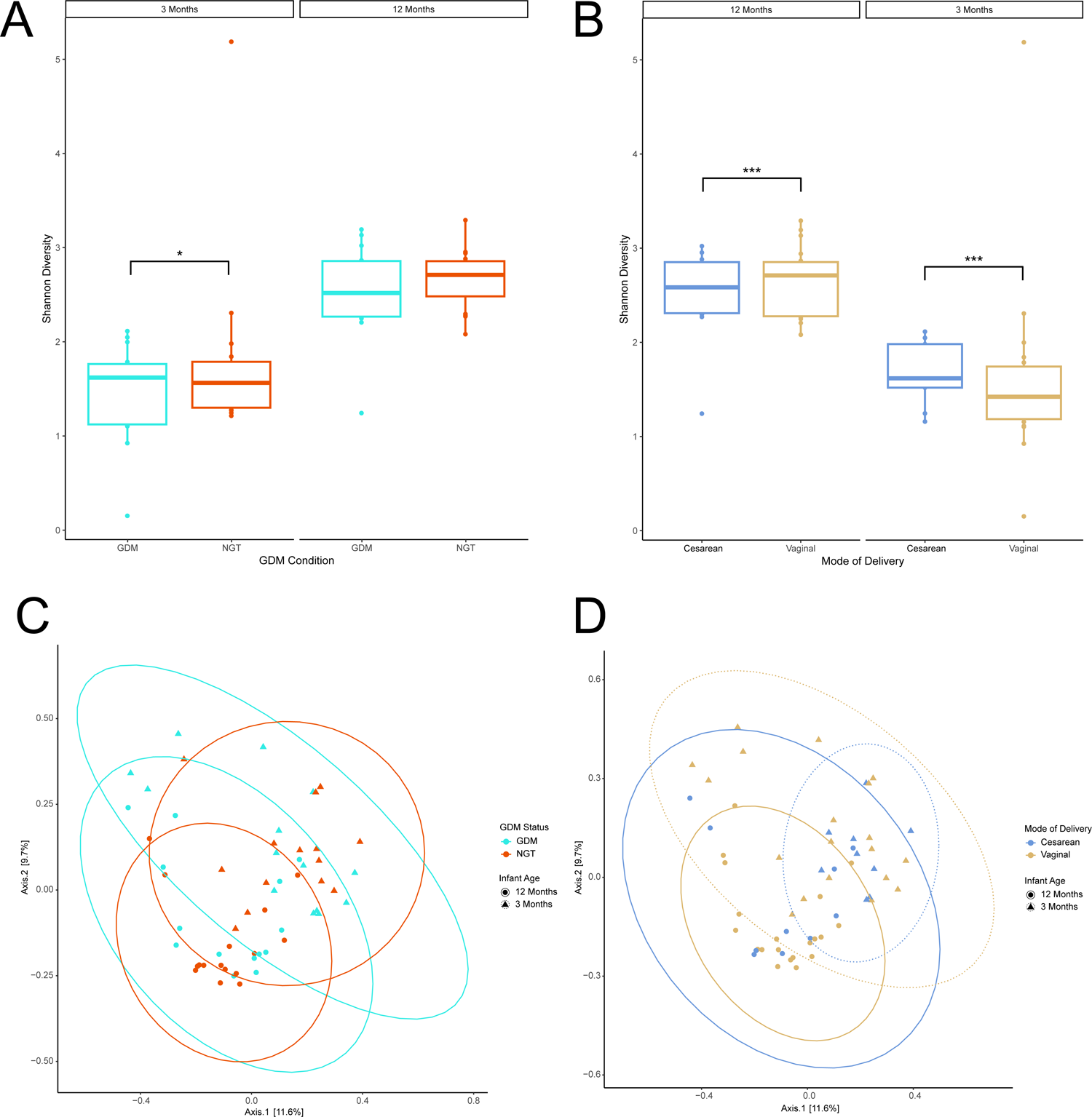
Diversity analyses of 16S data by mode of delivery. (a) Shannon Alpha diversity analysis. ANOVA was used to test for statistical significance. (b) Bray-Curtis PCoA Beta Diversity analysis. Ellipses represent 95% confidence interval for a multivariate normal distribution. n = 19, Cesarean-section; n = 39, Vaginal. n = 28, 3 months; n = 30, 12 months. * P ≤ 0.05, ** P ≤ 0.01, *** P ≤ 0.001.

Evaluation of beta-diversity (Supplementary Table 2) at the genus level through Bray-Curtis distances found that infant age (*P* < 0.001, PERMANOVA), mode of delivery (*P* = 0.045, PERMANOVA, Figure 2d), and exclusive breastfeeding (P = 0.004, PERMANOVA) had a significant effect on sample clustering. When examining beta diversity within 3-month samples, only exclusive breastfeeding status had a significant effect on clustering (*P* = 0.048, PERMANOVA). GDM status was not associated with any significant clustering effect (Figure 2c). Within 12-month samples, none of the comparisons tested had a significant clustering effect on beta diversity. These findings were also significant at the family level. At the OTU-level, only infant age (*P* < 0.001, PERMANOVA) and exclusive breastfeeding (P = 0.004, PERMANOVA) had significant effects on sample clustering. No significant effects were observed at the phylum level.

In order to uncover any taxa-level effects of GDM on gut composition, we examined individual changes in abundance of taxa (Supplementary Table 3). Using ANCOM-BC to test for differential abundance at a genus level, we found that 23 taxa were differentially abundant between the two infant ages, with 17 of these genera decreased in 12-month infants. These included *Enterococcus* (P = 3.61e-4, ANCOM-BC), *Klebsiella* (P = 4.29e-2, ANCOM-BC), and *Staphylococcus* (P = 4.96e-7), all decreased in abundance in 12-month-old infants. The only other comparison associated with differentially abundant taxa was mode of delivery at 12 months. Here, 2 genera were differentially abundant: *Fusobacterium* (P = 3.36e-2, ANCOM-BC), which was more abundant in c-section delivered infants, and *Streptococcus* (P = 9.85e-2, ANCOM-BC), which was more abundant in vaginally delivered infants. At a family level (Supplementary Table 4), Bacteroidaceae was decreased in C-section infants, across all infant ages (P = 3.23e-2, ANCOM-BC) and at 3 months (P = 4.65e-2, ANCOM-BC).

Overall, mode of delivery and infant age both had significant effects on alpha and beta diversity in several of the analyses we performed. Mode of delivery and infant age were both associated with differentially abundant taxa. Beta diversity analysis uncovered significant clustering by breastfeeding status across all infant ages and in the 3-month-old infants. No differentially abundant taxon nor significant clustering by beta-diversity was associated with GDM condition.

### Metatranscriptomics taxonomic expression differed significantly from 16S taxonomic composition

To functionally interrogate the gut microbiome of the 12-month-old infants, we performed metatranscriptomics on 30 unique samples, resulting in the generation of an average of 21154913.4 150bp paired-end sequence reads per sample (Table 2). After processing to filter low quality, host and rRNA, reads were annotated to genes and enzymes based on the ChocoPhlAn database, and taxonomy information retrieved using the MetaPro pipeline^21^. On average, 97.96% of reads were considered high quality and only an average of 0.10% of reads in each sample were considered host contaminants. Despite the use of the RiboZero Gold rRNA depletion kit, we still identified 53.00% of reads as either rRNA or tRNA. After filtering, an average of 44.68% of reads were retained, representing an average of 2979565.595 annotated putative mRNA reads.

**Table 2.**
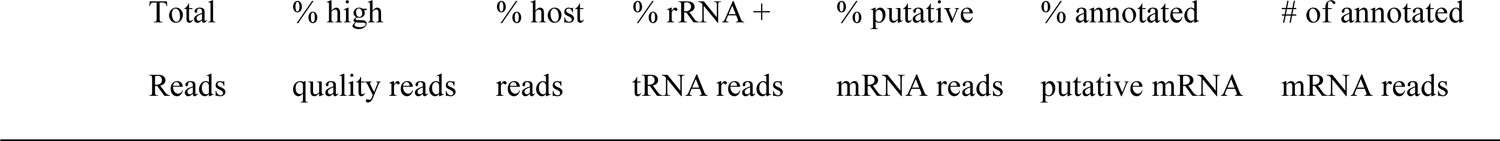
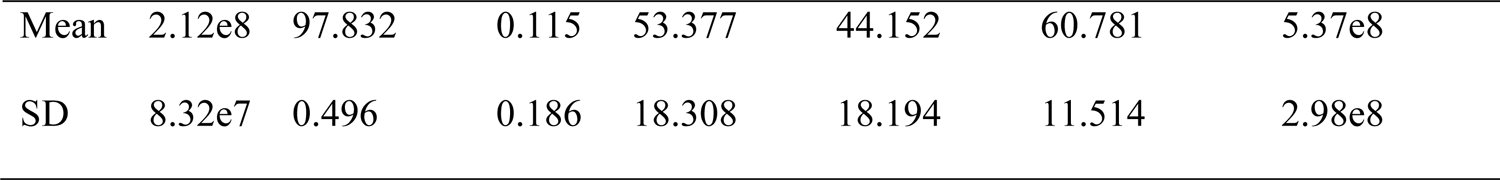
MetaPro Pipeline Metrics.

|  |  |  |  |  |  |  |
| --- | --- | --- | --- | --- | --- | --- |
| Total | % high | % host | % rRNA + | % putative | % annotated | # of annotated |
| Reads | quality reads | reads | tRNA reads | mRNA reads | putative mRNA | mRNA reads |

|  |  |  |  |  |  |  |  |
| --- | --- | --- | --- | --- | --- | --- | --- |
| Mean | 2.12e8 | 97.832 | 0.115 | 53.377 | 44.152 | 60.781 | 5.37e8 |
| SD | 8.32e7 | 0.496 | 0.186 | 18.308 | 18.194 | 11.514 | 2.98e8 |

Beta diversity analyses based on Bray-Curtis distances, found no single factor (GDM, mode of delivery, sex or exclusive breastfeeding status), or interaction between pairs of factors, associated with community structure in the metatranscriptomic datasets. Phylum rank was dominated by Bacteroidetes and Firmicutes in both the GDM and NGT samples (Supplemental Figure 1). DESeq2 analysis identified no differentially expressing taxa in the phylum rank by GDM condition. As with the 16S data, PERMANOVA indicated no significant differences in Bray-Curtis beta-diversity at neither the ranks of phylum (*P =* 0.378) or family (*P =* 0.741) between GDM and NGT samples.

In both GDM and NGT samples, the Bacteroidaceae family was most prevalent (Figure 3). Other prominent families include Ruminococcaceae, Lachnospiraceae, and Bifidobacteriaceae. To compare differences in individual taxon abundance and expression, we used DESeq2 to find families that exhibited a significantly different level of expression in metatranscriptomics samples. Of the 404 families detected here, 12 differentially expressing taxa were identified by DESeq2 between GDM and NGT infants. These include several high-abundance families such as Oscillospiraceae (stat = −3.33, *adjusted P* = 7.05e-3), Acidaminococcaceae (stat = 5.47, *adjusted P* = 2.46e-6), Rikenellaceae (stat = −5.14, *adjusted P* = 7.85e-6), Prevotellaceae (stat = 4.94, *adjusted P* = 1.46e-5), and Eubacteriaceae (stat = 3.26, *adjusted P* = 0.008). Additionally, Bray-Curtis clustering of gene expression was found to differ significantly from 16S taxonomic composition as found by PERMANOVA, further highlighting the difference between expression and composition. This discordance applied to the phylum rank (*P =* 9.99e-4), as well as the family rank (*P =* 9.99e-4).

**Figure 3.**
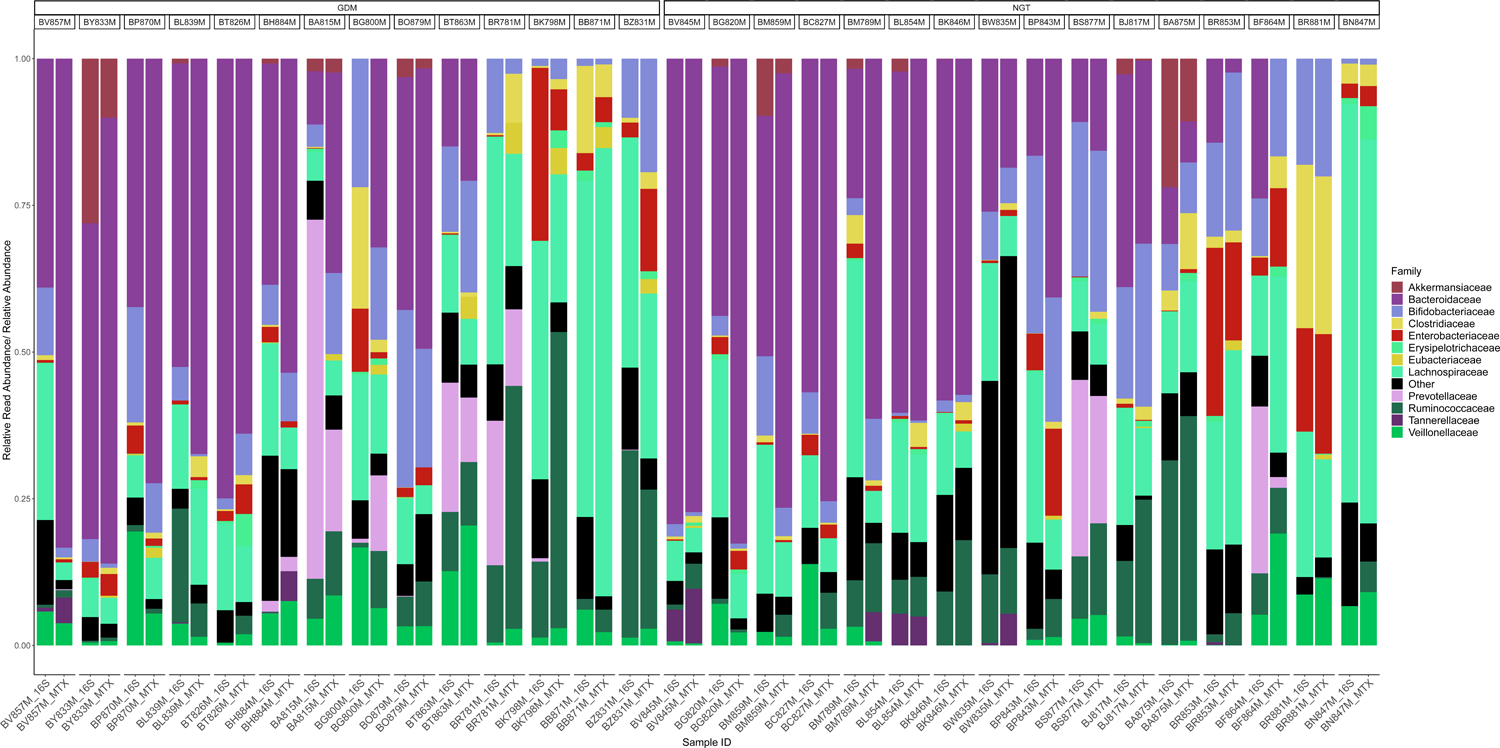
Stacked bar chart illustrating the relative abundance of families in the 16S samples compared to the relative read abundance expressed by each family in metatranscriptomics (MTX) samples. Samples are stratified by gestational diabetes (GDM) condition of the mother. Sorted by decreasing relative abundance of Bacteroidetes phylum in MTX samples. Limited to the top 12 most abundant families. Colours of each family correspond to colour of the parent phyla in Supplemental Figure 1 (e.g., Lachnospiraceae is colour coded to the Firmicutes phylum). n = 30.

### Mode of delivery has the largest impact on individual gene expression in the 12-month infant gut

Out of over 20,000 genes remaining post-filter, DESeq2 identified 294 differentially expressed genes (DEGs) associated with GDM status of the mother, 980 DEGs with mode of delivery, 260 DEGs with the sex of the infant, and 342 DEGs with EBF (Figure 4a, Supplemental Figure 2a-d, Supplemental Table 5). Considering the larger impact of mode of delivery on gene expression, each comparison was then re-examined after controlling for the impact of mode of delivery on function. Here, 226 genes were differentially expressed in the GDM status condition, 90 DEGs associated with sex, and 269 DEGs linked to EBF (Figure 4b-d).

**Figure 4.**
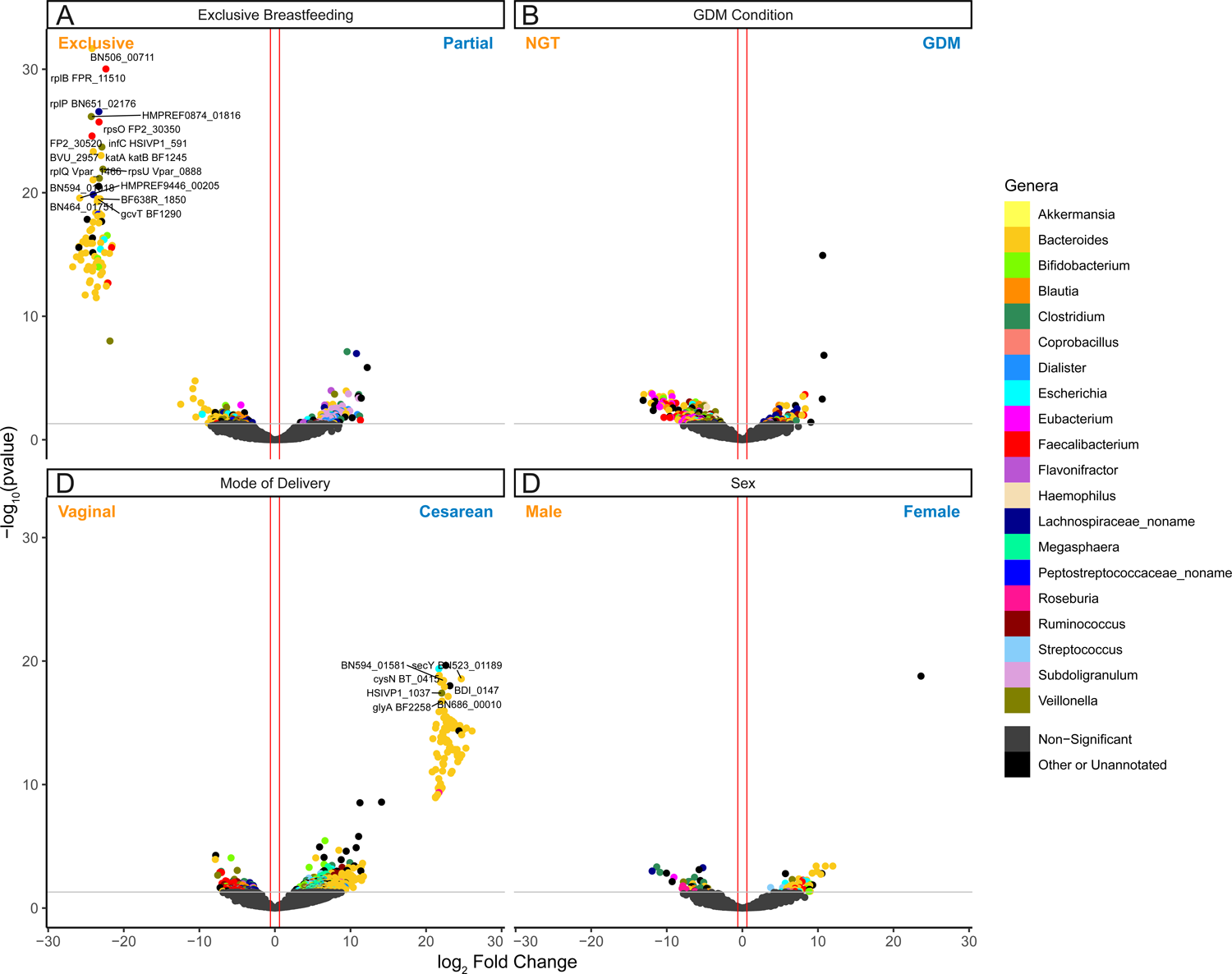
Volcano plot depicting DEG results for mode of delivery and mode of delivery-controlled factors. Log_2_ fold change and -log_10_(p value) are both from results of poscounts DESeq2. Blue points indicate DEGs that are upregulated in a condition. Red points indicate DEGs that are downregulated in a condition. Grey points are non-DEGs. (A) normal glucose tolerance (NGT) vs GDM. (B) Vaginal birth vs caesarean section birth. (C) Male vs female. (D) Exclusive breastfeeding vs. partial breastfeeding. Upregulation and downregulation are relative to the labels. n = 21563 genes.

Examination of the genera that express these DEGs reveals that *Bacteroides* are usually the most abundant taxa, with the exception of the mode of delivery-controlled GDM condition DEGs, where *Veillonella* expressed 20% of the genes. *Faecalibacterium*-annotated genes are also common across these DEGs. In the mode of delivery-controlled EBF DEGs, a significantly higher number of DEGs were annotated to *Subdoligranulum* than might be expected in the population (*P* < 0.001, hypergeometric test). *Subdoligranulum* was only otherwise present in the non-mode of delivery-controlled EBF DEGs. Despite being the second and third most common taxa respectively prior to differential expression, *Clostridium* and *Ruminococcus* were both significantly underrepresented in every DEG set apart from *Clostridium* in the mode of delivery-controlled sex DEG set (*Clostridium*: P < 1.25e-6; *Ruminococcus:* P < 0.0002 in all significant comparisons, hypergeometric test).

### Gestational diabetes condition was not associated with any enriched gene ontology terms

In order to examine which broader categories of function were represented by these DEGs, we performed a gene set enrichment analysis using the fgsea R package^22^. Here, we examined enrichment using three different databases: the biological process ontology of the gene ontology resource (GO; Figure 5a), the comprehensive antibiotic resistance database (CARD; Supplemental Figure 3), and the carbohydrate-active enzymes database (CAZy/ CAZyme; Figure 5b)^23–25^. These resources were chosen to categorize function on a higher level, and to specifically observe how function might change in an area of interest, notably microbial resistance to antibiotics and differences in carbohydrate metabolism. Within the single factor comparisons, only sex and mode of delivery DEGs were associated with any enriched terms across all three databases. Considering the outsized number of DEGs associated with mode of delivery, we decided to examine set enrichment primarily within the mode of delivery-controlled set of DEGs for sex, GDM status, and exclusive breastfeeding.

**Figure 5.**
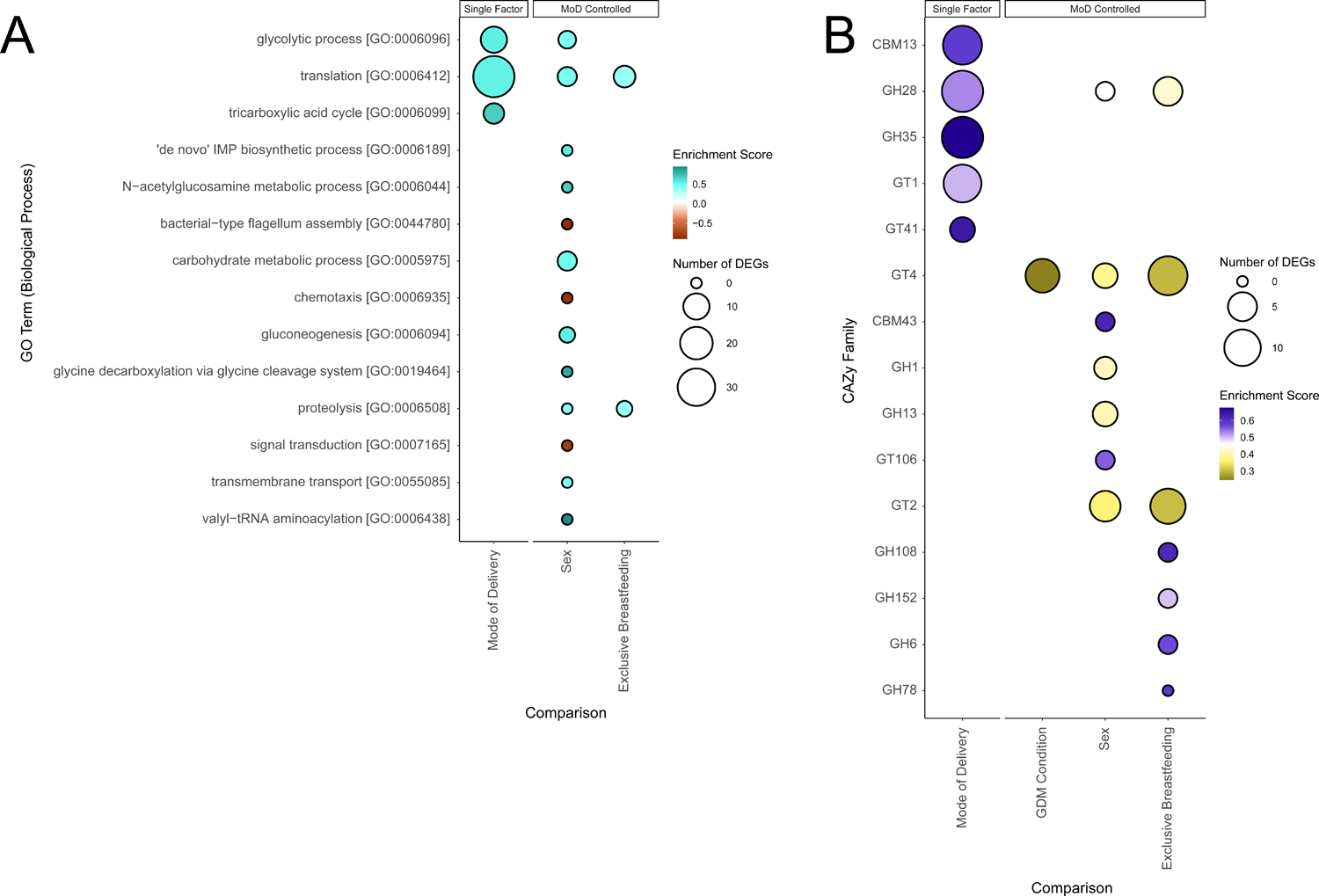
Bubble charts detailing each Gene Ontology (GO) biological process term and carbohydrate-active enzyme (CAZy) family that was significantly enriched in our differential expression results. Enriched terms were identified using the fgsea R package^22^. Size of bubble indicates the number of DEGs from a given functional category associated with the factor. Enrichment score indicates degree and direction of enrichment for a given term. GO terms and CAZy families are sorted by adjusted p-value. Mode of delivery (MoD) controlled terms are indicated. (A) Bubble chart of GO term enrichment. MoD-controlled GDM condition is omitted as there were no significant terms. (B) Bubble chart of CAZy enzymes. CBM = Carbohydrate-Binding Modules, GH = Glycoside Hydrolases, GT = GlycosylTransferases.

Mode of delivery was associated with an increase of tricarboxylic acid cycle (TCA; P 2.46e-2, fgsea), glycolytic process (P = 2.46e-2, fgsea), and translation (P < 0.001, fgsea) GO terms in C-section infants. Similarly, in the CAZy enrichment, mode of delivery was associated with 5 CAZy families, all of which upregulated in C-section infants as well. Mode of delivery was also the only single factor to display any enrichment in the CARD database, including an increase in tetracycline (P = 2.32e-2, fgsea) and isoniazid resistance (P = 2.26e-2, fgsea), and several antibiotic efflux pump terms in C-section infants, and a decrease in a vancomycin resistance term (P = 2.32e-2, fgsea).

Neither single factor GDM nor EBF DEGs were associated with any enriched sets in any of the three databases. However, controlling for mode of delivery, GDM DEGs were enriched for two functional terms: the glycosyl transferase (GT) family 4 CAZyme (P = 9.03e-4, fgsea), and the fusidic acid resistance CARD resistance family (P = 3.97e-2, fgsea), both enriched within GDM infants. EBF DEGs were enriched for 7 CAZy families, including 5 glycoside hydrolase (GH) families, and two CARD sets, all of which are increased in enrichment in partially breastfed infants, apart from an isoniazid resistance term (P = 4.26e-2, fgsea). The significant GO terms here include translation (P = 5.88e-3, fgsea) and proteolysis (P = 3.58e-7, fgsea), both of which are increased in partially breastfed infants.

### Applying alternative differential expression techniques to metatranscriptomics data

In addition to DESeq2, we attempted several other methods for differential expression analysis. These methods include those designed originally for use with 16S rRNA datasets, such as ANCOM-BC^26^ and ALDEx2^27^, and one designed for metatranscriptomics, taxon-specific scaling^28^. Both ANCOM-BC and ALDEx2 address the compositional nature of microbiome data, and have been demonstrated by previous benchmarking studies to be stringent in their determination of differential abundance^29^. As such, both ANCOM-BC and ALDEx2 produced no DEGs across all four conditions, GDM status, mode of delivery, sex, and EBF status. Additionally, we explored the use of consensus DEGs between both DESeq2 and a second differential analysis tool, corncob^30^. While this increased the likelihood of a given DEG to be a true positive, this method only identifies DEGs without the gene-level statistics required for downstream functional enrichment analyses.

Taxon-specific scaling is a normalization method that uses the DESeq2 framework to resolve an issue with metatranscriptomics data where changes in the abundance of a taxon expressing a gene could be mistaken for changes in expression of that gene^28^. Overall, this method produced fewer DEGs compared to DESeq2. GDM condition was associated with 93 DEGs, while 171, 59, and 109 DEGs were assigned to mode of delivery, sex, and EBF, respectively. However, fgsea analysis revealed 6 GO terms enriched in the GDM infants: DNA topological change, glycolytic process, mRNA catabolic process, protein folding, protein refolding, and translation. Mode of delivery yielded only 1 GO term (translation, P = 1.76e-6, fgsea), sex yielded no terms, and exclusive breastfeeding status produced two terms - fatty acid biosynthetic process (P = 4.57e-2, fgsea) and protein refolding (P = 4.57e-2, fgsea).

## DISCUSSION

This case control study sought to profile the changes experienced by the infant gut microbiome in response to a pregnancy complicated by GDM, while concurrently exploring the effect of mode of delivery, breastfeeding, and sex. While previous studies have examined the impact of GDM on the composition of the infant gut microbiome, we believe this is the first study that incorporates a functional metatranscriptomics approach. Our results indicate that maternal GDM is associated with a decrease in alpha diversity across all infant ages and in the 3-month infants specifically. This corroborates existing abundance-based microbiome literature, in which alpha diversity has been shown to be significantly lower in neonates of mothers with GDM^31–33^. This is not always consistent in literature; Crusell et al. found no differences in Shannon’s diversity nor Pielou’s evenness indices at either 1 week nor 9 month infant ages^14^. However, there was a decrease in OTU richness at 1 week that was not observable at 9 months. Koren et al. additionally found no difference in species richness between GDM exposed and NGT exposed children^34^. As GDM did not have a significant impact on alpha diversity in the 12-month infants, it is possible that alpha diversity in GDM infants increases to meet the levels of NGT infants sometime before 12 months of age. Infants are exposed to a range of environments during the first months of life which may act to mitigate the effects of GDM on early microbial composition^3^ and increase alpha diversity.

Our data did not show a statistically significant difference in Bray-Curtis distances between GDM and NGT samples across both infant ages, suggesting that the gut microbiome compositions of GDM infants in our study were not significantly more closely related to each other than to NGT infants. The effect of GDM here could perhaps be masked by other stronger factors. Previous literature indeed have found differences in beta diversity in GDM infants^13,14,33,34^. Koren et al. demonstrated that differing beta diversity may slowly return to normal by 4 years of age^34^. The infant microbiome approaches a close resemblance with an adult gut composition around 3 years of life, at which point these differences in beta diversity would likely have resolved^3^. In our data, it is possible that the NGT mothers having already failed the GCT meant that the interindividual differences between GDM and NGT infant guts were not distinct enough.

In addition to the GDM status of the mother, we also examined the impact of several other factors on alpha and beta diversity. Mode of delivery was linked with a significant difference in alpha diversity in 3-month and 12-month infants, though not when testing across all infant ages. Mode of delivery also significantly affected beta diversity across all 16S timepoints. Existing literature offers somewhat conflicting results in regards to how mode of delivery induces compositional changes to the microbiome over the first year of life^35–39^. While mode of delivery has a considerable impact on the gut microbiome early on, some studies suggest its influence wanes by 6 months of age, possibly as early as 8 weeks^39–43^. Rutayisire et al. conducted a review of 7 studies that found lower diversity within the Bacteroidetes and Actinobacteria phyla along with a lower abundance of *Bacteroides* associated with a C-section birth within the first 90 days postpartum^42^. After this time period, the influence of mode of delivery seemed to wane, as studies found no difference in *Bacteroides* colonization by 6 months. Previous research centred on the gut microbiome of older children (≥ 3 years) have found few differences in overall diversity in the microbiome associated with birthing method, though individual taxa had changed, including the *Clostridium* genus^44,45^. Our results support the notion that perturbations to alpha and beta diversity by delivery remain detectable within the first year.

In our 16S samples, we found a reduction of Bacteroidaceae in C-section infants across both infant ages and at 3 months specifically. This is a well-recorded phenomenon, due to C-section infants receiving less exposure to the maternal microbiome^42,46^. In comparison, 16 families were differentially expressive in the metatranscriptomics samples, which also include a reduction in Lactobacillaceae and Streptococcaceae transcripts in C-section delivered infants. Both of these families are lactic acid bacteria, and the former contains many species considered beneficial to the infant gut, such as *Lactobacillus acidophilus* and *Lacticaseibacillus rhamnosus*^47–50^. These taxa were likely passed down to the vaginally delivered infants from the vaginal canal^51^, and the reduction in abundance of these species could represent a long-term consequence of C-section delivery. A longitudinal metagenomics study performed by Bokulich et al. suggests that mode of delivery has a significant impact on diversity measures and taxa abundance during the first two years of life, but these effects decrease in size as the infant microbiome begins to reach an adult-like state^36^. Our results support these findings, though future longitudinal studies may elucidate whether differences in infant health correlate with gut microbiome composition.

Since there were no mothers in this study that exclusively used infant formula milk, groups compared between exclusive and partial breastfeeding diets. Therefore, the effect being measured is likely more subtle than comparison between exclusively breastfed and exclusively formula-fed infant gut microbiota. Breast milk aids in nurturing infant gut microbes through a supply of human milk oligosaccharides (HMOs) among other metabolites, vitamins and immunoglobulins^48,52^. These HMOs are utilized by *Bifidobacterium* and *Bacteroides* species in the gut^53^, the latter of which make up a large proportion of transcripts in our data. A meta-analysis of 7 studies conducted by Ho et al. found that non-exclusively breastfed infants under 6 months of age displayed higher gut Shannon alpha diversity^54^. Breastfeeding status also had a strong effect on our 16S data, with partially breastfed infants experiencing a decrease in alpha diversity across all infant ages and at 3 months. Similarly, breastfeeding status was associated with significant clustering by beta-diversity at all infant ages and at 3 months. Once again it is possible these differences in diversity are eventually lost by 12 months, as a study of 423 Finnish adolescents found no differences in alpha nor beta diversity depending on early breastfeeding diet^55^.

The discrepancy in relative abundance of taxas between the 16S rDNA and metatranscriptomics datasets likely reflects differences associated with species abundance and their transcriptional activity. While we found several families to differ in expression in the metatranscriptomics data, 16S analyses did not uncover any differentially abundant taxa at any level. This may also be explained by the use of ANCOM-BC, a tool built for 16S analysis, disagreeing with DESeq2^26^.

Few studies on the infant gut microbiome have applied metatranscriptomics to study the infant gut microbiome, and they have tended to focus on preterm births^56–58^. Beyond these, one study reported pathway changes in a longitudinal study over the first year of life^59^, while another study profiled changes in microbiome activity alongside solid food adaptation, outlining steps towards an adult-like composition^60^. In comparison, our metatranscriptomics results focused on twelve-month old infants, aiming to reveal factors that may have sustained effects on microbiome activity. Taking into account both the number of DEGs associated with these factors and the number of molecular function GO terms enriched within those DEGs, it is likely that by 12 months, the persisting effect of maternal GDM status and EBF is modest, while mode of delivery and sex both influence the function of the microbiome.

As there have been no metatranscriptomics studies on GDM, most predictions on the functional changes GDM may induce are derived from correlating taxonomic abundance changes to factors in health^32,61^. For example, one study found a positive correlation between maternal oligosaccharide consumption and abundance of *Ruminococcus*, linking this with an increase to butyrate and bacteriocin production that is associated with this taxa^32^. Here we applied metatranscriptomics to more directly assess functional changes within the microbiome. However, in our data, we found no significantly enriched functions, as defined by GO terms, even after controlling for the overarching effects of mode of delivery. Though there were 6 GO terms enriched in GDM infants when examining the taxon-specific scaling results, there was only one term related to carbohydrate metabolism, the rest being non-specific growth-related terms. We found only one significantly enriched CAZy family (GT4) upregulated in GDM infants after controlling for mode of delivery, despite GDM being primarily an issue of carbohydrate metabolism in the host. However, GTs have previously been implicated in diabetes as a potential therapeutic target^62^ and have been observed to change in activity in response to hyperglycemia^63^. These results suggest that, by 12 months of age, there is no significant persistent perturbation of functional categories in the infant gut resulting from maternal GDM status. Any prevailing effect might exist as a slight increase in growth or carbohydrate metabolism in the GDM infant gut. As highlighted below, we cannot rule out the possibility that the diet of the mothers diagnosed with GDM in this study were able to overcome any potential negative effects of GDM on the infant gut, which has previously been shown through metagenomics by Sugino et al.^64^. The study compared a conventional diet with a diet that was higher in complex carbohydrates and lower in fat in women diagnosed with GDM. As a result, infants of the mothers fed the treatment diet displayed increased gut alpha diversity over time. Indeed, previous work from our cohort has consistently demonstrated lower birthweight of infants born to women with GDM, compared to those with normal or milder dysglycemia during pregnancy, which indicates excellent glycemic control in this GDM cohort during pregnancy^65^.

Three terms were significantly enriched in mode of delivery: translation, glycolytic process, and tricarboxylic acid cycle. Translation correlates with cell growth, potentially signifying an increase in overall microbial growth in C-section infants within this study^66^. The other two carbohydrate metabolism-related terms are also upregulated in infants delivered through C-section, which, together with translation, suggests that there is increased microbial cell proliferation in infants delivered through C-section compared to vaginally delivered infants. This is supported by the overall direction of expression for most mode of delivery DEGs (Fig. 3b) and the direction of the significantly enriched CAZy families (Fig. 4b), all of which are enriched in samples associated with C-sections. Interestingly, this is somewhat contrary to analyses of the early microbiome, in which vaginally delivered infants tend to be associated with a higher diversity and enrichment of metabolic processes^67,68^. Along with the high number of DEGs and the observed changes in diversity, our results point to mode of delivery as having a clear effect on both the structure and function of the 12-month-old infant gut microbiome. As previously mentioned, it is unclear whether mode of delivery still affects infant gut microbial composition at 12 months of age^35–39^. While there are few studies that examine the impact of mode of delivery on function, these primarily concern the impact that C-section birth might have on long-term immune development^69,70^. Overall, we found mode of delivery has a strong effect on infant gut microbial diversity and function, even at twelve months after birth.

Post mode of delivery-control, sex was associated with 13 enriched GO terms, the most of any comparison tested. It should be noted however that many of these pathways did not contain DEGs identified by DESeq2, indicating a functional category-wide enrichment that was not driven by any particular DEGs. Five terms were directly related to nutrient metabolism, primarily carbohydrate metabolism, all of which had been enriched within female infant gut microbiomes. Similarly, seven CAZy families were significantly enriched, though these were not conformally unidirectional. The remaining GO terms are relatively unspecific terms related to translation or motility, with both motility terms (eg, chemotaxis, bacterial-type flagellum assembly) enriched in the male samples. PICRUSt analysis of 16S data has previously identified some sex-specific differences between pre-term, dizygotic twin male and female gut function; though there were carbohydrate metabolism pathways differentially regulated in these results, many other unrelated pathways were differentially regulated as well^71^. There have been previous studies in adult gut microbiomes studying sex-specific differences in carbohydrate metabolism, but the mechanism driving this requires further study^72,73^. Hormones appear to affect the composition of the infant gut since birth^74^, and our results point to some early microbiome differences in function as well, relating to metabolism and other core bacterial processes.

Only two GO terms – proteolysis and translation – were enriched in the mode of delivery-controlled EBF. A previous meta-analysis by Ho et al.^54^ involving infants less than six months of age found that most KEGG pathways that differed between partially and exclusively breastfed infants were involved with metabolism. Metabolomic analysis of four-month-old infant fecal samples found that breastfeeding status affects the composition of the metabolites produced in the gut^75^. Though in our data there were seven CAZy families significantly enriched between infants of varying EBF status, they did not neatly fall into one direction or the other. Once again, the differences observed between our data and other studies might be explained by age differences – many of the studies examine the infant gut between 4-6 months of age, and the infant diet may change greatly in the subsequent six months.

Metatranscriptomics data can be difficult to analyze due to sparsity, compositionality, and the lack of dedicated tools built solely for its intricacies^76,77^. Consensus with a second differential expression method or implementation of taxon-specific scaling may offer a safer alternative, though this makes downstream analyses like gene set enrichment difficult. Notably, Zhang et al.^76^ iterated upon the original method of taxon-specific scaling^28^ with the additional use of metagenomic data in the normalization step, although we did not implement this method as we did not perform metagenomic sequencing in our study. ANCOM-BC and ALDEx2 have been shown to exhibit stringent false positive detection in previous benchmarking studies at the cost of lower statistical power^29,78^. In our study, neither of these methods returned any DEGs, though it is difficult to determine whether this is a result of low true positives or their incompatibility with metatranscriptomics data.

Two additional methodological considerations in this study warrant further discussion. Firstly, the infants of mothers with GDM were monitored during pregnancy in a tertiary care centre, and their infants demonstrate lower birthweight than the control group, suggesting effective glucose management during pregnancy^65^. As a result, this may have acted to mitigate the potential microbiome-modifying effects of the condition. Secondly, the reference control group was composed of mothers who failed the initial GCT but did not meet criteria for GDM on glucose tolerance testing. While glycaemic management intervention is therefore not indicated for this group of mothers, they are experiencing subclinical insulin resistance. Therefore, we must consider the possibility that even exposure to insulin resistance may alter the infant gut microbiome in a manner which is similar to GDM.

While GDM has been observed to affect the composition of the early infant microbiome, in this study we were able to elucidate the functional consequences of GDM further into the development of the infant gut. Though our results support some changes in beta diversity, GDM exerted little effect on function in the 12-month-old infant gut. As the gut develops with the introduction of solid foods and weaning from breastfeeding, the effect of GDM exposure becomes less prominent against an increasingly complex gut community. Similarly, while there may be differences in diversity remaining, our results suggest that the impact of EBF wanes by one year. Some functional differences are visible at 12 months in sex and mode of delivery, with the latter likely having the greatest influence. These data serve to reassure parents and paediatricians that, while mode of delivery appears to impact function and diversity for longer than anticipated, GDM may not have persistent effects on the function nor composition of the infant gut microbiome, at least when compared to mothers who did not experience GDM. Regulation of diet and glycemic control may alleviate the potential impact of GDM on the infant gut microbiome. Future studies in this area could explore the mitigating effects of diet and other interventions on the impact of GDM on the infant gut. Future work may also consider the use of paired metagenomic or metabolomic data to improve metatranscriptomics differential expression analysis and to build a robust profile of how expression might correlate with copy number or nutrient availability.

## METHODS

### Study Design

Pregnant persons with no prior diagnosis of diabetes were recruited in late 2^nd^ to early 3^rd^ trimester at Mount Sinai Hospital, Toronto. The cohort was enriched for those that failed a 1h 50g glucose challenge test. At 29.9 ± 2.9 weeks gestation, participants underwent a 2-h 75g oral glucose tolerance test to determine GDM status^9^.

Participating women attended study visits that included nurse-administered questionnaires for personal medical, obstetrical, and family history, anthropometric measurement, and tests of lipid profile, insulin sensitivity, glucose tolerance, and adipokines in late pregnancy and at 3-months and 12-months postpartum. Offspring were assessed at birth, 3, and 12 months. Infants were assessed for anthropometric measurements, and the parent completed questionnaires related to their infant’s health and nutrition. At 3-month and 12-month visits, stool samples were collected for assessment of gut microflora. Metatranscriptomic analyses were performed on a subset of 12-month samples to compare mothers with normal glucose tolerance to those with gestational diabetes.

The exclusion criteria for this study were as follows:

– Infants born less than 37 weeks gestation or greater than 42 weeks of gestation
– Multiparity
– Infants with significant medical illness requiring prolonged or repeated hospitalization
– Infants with medical conditions or taking medications (e.g. glucocorticoids) known to alter cardiometabolic risk
– Stool samples of infants who had antibiotics and/or probiotics within 3 months of collection were also excluded from microbiome analysis

Data was collected from the chart and by questionnaires for the following:

– Parental BMI
– Maternal pre-gravid body mass index
– Weight gain during pregnancy
– Mode of delivery
– Infant feeding history (exclusive breastmilk vs formula and breast milk; age of introductions of solids)
– Infant Sex
– Infant Birth Weight
– Gestational Age at Birth
– Other factors that may affect microbiome (ex. Antibiotic, probiotic use)

### 16S Sample Preparation and Analysis

DNA and RNA stool samples were homogenized prior to extraction. DNA extraction was completed using Omega E.Z.N.A.TM Stool DNA Isolation Kit, and DNA quality was assessed spectrophotometrically. High-throughput sequencing of the hypervariable V4 region of the 16S rRNA gene, using 150 bp primers in PCR, was performed at The Centre for the Analysis of Genome Evolution and Function (CAGEF) at the University of Toronto. Primers were adapted to incorporate Illumina adapters with indexing barcodes and sequenced using MiSeq. QIIME 2 v2023.2 was used to process raw FASTQ files, remove low-quality reads, and assemble and merge paired reads^79^. Deblur v1.1.1 was used to cluster reads into ASVs^80^. Samples were visualized using phyloseq 1.44.0 and ggplot 3.4.1^81,82^. Adonis2 from the vegan 2.6-4 R package was used to conduct PERMANOVA statistical tests on beta diversity and ANCOM-BC 2.1.2 was used to test for differential abundance^26,83,84^. ANOVA was used to assess for differences in alpha diversity. Notably, 2 infants did not have 3-month data, and 1 did not have breastfeeding information for the 3-month timepoint.

### Metatranscriptomics Sequencing

Stool collected at 12-months was processed using the RNeasy PowerSoil Total RNA kit, using 400-500 mg of each sample. Changes to the kit protocol include an incubation with SR4 at 1 hour at room temperature at step 9, heat pellet at 45°C for 10 minutes at step 12, incubation with SR4 overnight at −20°C at step 17, and elution in 100 μL of SR7. DNA was removed from the sample by incubating 44 μL of RNA with 1 μL of Turbo DNase and 5 μL of Turbo 10X Buffer at 37°C for 30 minutes. RNA was cleaned with Zymo RNA clean-up column and processed by BioAnalyze for quality and quantity of RNA. 2.5 μg was added to RiboZero Gold kit to remove rRNA^85^. cDNA was amplified using Ultra directional RNA kit, following protocol for Ribosome Depleted RNA. Qubit/ PicoGreen was applied to the barcoded cDNA libraries, and the final library was processed by BioAnalyze on DNA chip to determine size. Diluted to 4 nM, denatured and loaded on NextSeq high output 150bp x 2 sequencing kit at 1.9 pM with 1% PhiX spike-in as standard.

### Metatranscriptomics Reads Processing

Reads were processed to remove low-quality reads and other unnecessary reads using the MetaPro pipeline, which also annotates gene reads using the ChocoPhlAn pangenome database from HUMAnN 2.0 and NCBI non-redundant (NR) protein databases^86–88^. Enzyme annotations are assigned according to the Swiss-Prot database and taxonomic annotations also rely on ChocoPhlAn and NCBI NR database, in addition to Kaiju 1.9.0 and Centrifuge 1.0.4^89–91^. The pipeline begins by removing low-quality reads, adapter sequences, and duplicate reads. Low-quality reads are defined as having a quality score below 75 according to FastQC 0.11.9^21^. Adapter sequences are identified and removed using AdapterRemoval v2.1.7 in tandem with VSEARCH v2.7.1 ^92,93^. In the vector contamination removal step, pBLAT 2.0 and BWA 0.7.17 were used to search reads for matches with contaminants and the UniVec_Core dataset was also utilized for its library of known vector and sequencing contaminants^88,94,95^. Host reads corresponding to the GRCh38 human reference genome were removed next, followed by use of BARRNAP v0.9 and Infernal to remove abundant rRNA sequences^96,97^. Duplicate reads that were previously removed are added back in, and the mRNA is then assembled into contigs using the rnaSPAdes v3.14.1 transcript assembly algorithm and MetaGeneMark v1 to annotate contigs to putative genes^98,99^. Contigs are then annotated to genes and assigned taxonomic information before annotated with enzyme function. Metapro utilizes three sequence similarity search tools in tiers, starting with the least sensitive and increasing in sensitivity^86^. In sequence, these tools are BWA 0.7.17, pBLAT 2.0, and DIAMOND 0.9.19^94,95,100^. Enzyme annotation is carried out using DETECT v2, PRIAM version 2018, and DIAMOND 0.9.19 ^100–102^. Finally, annotated outputs are generated for use in bioinformatic analysis, and the results are visualized in Cytoscape v3.9.1 as a KEGG pathway^103,104^.

### Metatranscriptomics Analyses

adonis from the vegan 2.3-5 R package was used to perform analysis of variance using distance metrics and incorporating multiple factors to compare Chao1 alpha diversity between samples, and Bray-Curtis and Weighted and Unweighted UniFrac beta diversity between samples^84^. Data for differential expression was filtered to remove genes with fewer than 6 counts in 6 samples, resulting in 21563 genes for differential expression and set enrichment analysis. DESeq2 3.16 with the poscounts size factor estimator was used to identify differentially abundant taxa, differentially expressed genes, and differentially expressed enzymes^105^. Taxon-specific scaling was adapted from a technique described by Klingenberg and Meinicke^28^, with changes made to data manipulation portions of the original code to fit the structure of our data. Other differential expression techniques we attempted include ANCOM-BC 2.1.2, ALDEx2 1.30.0, and corncob 0.3.1^26,27,30^. GO term, CAZy, and CARD gene set enrichment analysis was done using fgsea 1.26.0^22^. GO terms were retrieved using UniPro ID mapper^89^. CAZy annotations were retrieved using dbCAN3^106^. CARD annotations were retrieved using CARD’s Resistance Gene Identifier^24^. Visualizations of figures 2-5 and supplemental figures 1-3 were done through ggplot 3.4.1^82^.

### Ethics statement

The research ethics board at The Hospital for Sick Children and Mount Sinai Hospital approved the protocol. All mothers provided written informed consent for their child. The work described was carried out in accordance with the code of ethics of the World Medical Association (Declaration of Helsinki) for experiments involving humans. Informed consent was obtained from all research participants according to the protocol approved by the Research Ethics Board of the Hospital for Sick Children and Mount Sinai Hospital (REB# 1000007561).

### Author contributions

JH and JP conceived and designed the study. KH organized patient recruitment and collection of samples and metadata. PWW and DG supervised extraction of nucleic material for high throughput sequencing. RVC and JC performed sequence data analyses. RVC, JP, JH, and PR wrote the manuscript. All authors reviewed and/or edited the paper.

## Acknowledgements

This work was funded by the Heart & Stroke Foundation of Canada (G-14-0006308) to JH and JP; and the Canadian Institutes for Health Research (MRT-168043) to JP. Computing resources were provided by the SciNet High Performance Computing (HPC) Consortium; SciNet is funded by the Canada Foundation for Innovation under the auspices of Compute Canada, the Government of Ontario, Ontario Research Fund - Research Excellence, and the University of Toronto. We would like to thank to Rebecca Noseworthy for her assistance and feedback on the manuscript.

## Data availability

Sequence data is deposited at the NCBI under BioProject repository identifier PRJNA1013505.

## TABLES AND FIGURE LEGENDS

**Supplementary Table 1.** Study Demographic Table by Infant Age and Data.

|  |  | 16S |  | Metatranscriptomics |
| --- | --- | --- | --- | --- |
|  |  | 3 Months | 12 Months | 12 Months |
| GDM Condition | GDM | 14 | 14 | 14 |
|  | NGT | 14 | 16 | 16 |
| Mode of Delivery | Vaginal | 19 | 20 | 20 |
|  | Caesarean Section | 9 | 10 | 10 |
| Sex | Female | 11 | 11 | 11 |
|  | Male | 17 | 19 | 19 |
| EBF | Exclusive | 11* | 8 | 8 |
|  | Partial | 16 | 22 | 22 |
| Total Samples |  | 28 | 30 | 30 |

\* One 3-month sample did not record breastfeeding status

**Supplementary Table 2.** Beta-Diversity PERMANOVA Results.

|  | All Infant Ages<br>(p-values) | 3 Months (p-values) | 12 Months<br>(p-values) |
| --- | --- | --- | --- |
| Visit | 0.001 | - | - |
| GDM Condition | 0.713 | 0.991 | 0.559 |
| Mode of Delivery | 0.069 | 0.105 | 0.485 |
| Sex | 0.695 | 0.952 | 0.322 |
| EBF | 0.008 | 0.039 | 0.634 |
| Gestational Age | 0.390 | 0.369 | 0.618 |
| Infant Birth BMI-z | 0.403 | 0.341 | 0.424 |

**Supplementary Table 3.** ANCOM-BC Number of Differentially Abundant Genera.

|  | All Infant Ages | 3 Months | 12 Months |
| --- | --- | --- | --- |
| Visit | 25 | - | - |
| GDM Condition | 0 | 0 | 0 |
| Mode of Delivery | 0 | 0 | 2 |
| Sex | 0 | 0 | 0 |
| EBF | 0 | 0 | 0 |

**Supplementary Table 4.** ANCOM-BC Number of Differentially Abundant Families.

|  | All Infant Ages | 3 Months | 12 Months |
| --- | --- | --- | --- |
| Visit | 11 | - | - |
| GDM Condition | 0 | 0 | 0 |
| Mode of Delivery | 1 | 1 | 0 |
| Sex | 0 | 0 | 0 |
| EBF | 0 | 0 | 0 |

**Supplementary Table 5.** DESeq2 Number of Differentially Expressed Genes.

|  | Single Factor | Mode of Delivery Controlled |
| --- | --- | --- |
| GDM Condition | 294 | 226 |
| Mode of Delivery | 980 | - |
| Sex | 260 | 90 |
| EBF | 342 | 269 |

### Figure Legends

**Supplemental Figure 1.** Stacked bar chart illustrating the relative abundance of phyla in the 16S samples compared to the relative read abundance expressed by each family in metatranscriptomics (MTX) samples. Samples are stratified by gestational diabetes (GDM) condition of the mother. Sorted by decreasing relative abundance of Bacteroidetes phylum in MTX samples. Limited to the top 7 most abundant phyla. Colours of each phylum correspond to colour of the child family in Figure 3. n = 30.

**Supplemental Figure 2.** Volcano plot depicting differentially expressed gene (DEG) results for single factors. Log_2_ fold change and -log_10_(p value) are both from results of poscounts DESeq2. Blue points indicate DEGs that are upregulated in a condition. Red points indicate DEGs that are downregulated in a condition. Grey points are non-DEGs. (A) Exclusive breastfeeding vs. partial breastfeeding. (B) NGT vs GDM. (C) Vaginal birth vs caesarean section birth. (D) Male vs female. Upregulation and downregulation are relative to the labels. n = 21563 genes.

**Supplemental Figure 3.** Bubble charts detailing each Comprehensive Antibiotic Resistance Database (CARD) resistance gene family that was significantly enriched in our differential expression results. Enriched terms were identified using the fgsea R package^22^. Size of bubble indicates the number of DEGs from a given gene family associated with the factor. Enrichment score indicates degree and direction of enrichment for a given term. CARD resistance genes are sorted by adjusted p-value.

