## Supplementary figures and images for "The impact of gestational diabetes on functional capacity of the infant gut microbiome is modest and transient"

### Supplemental Figures

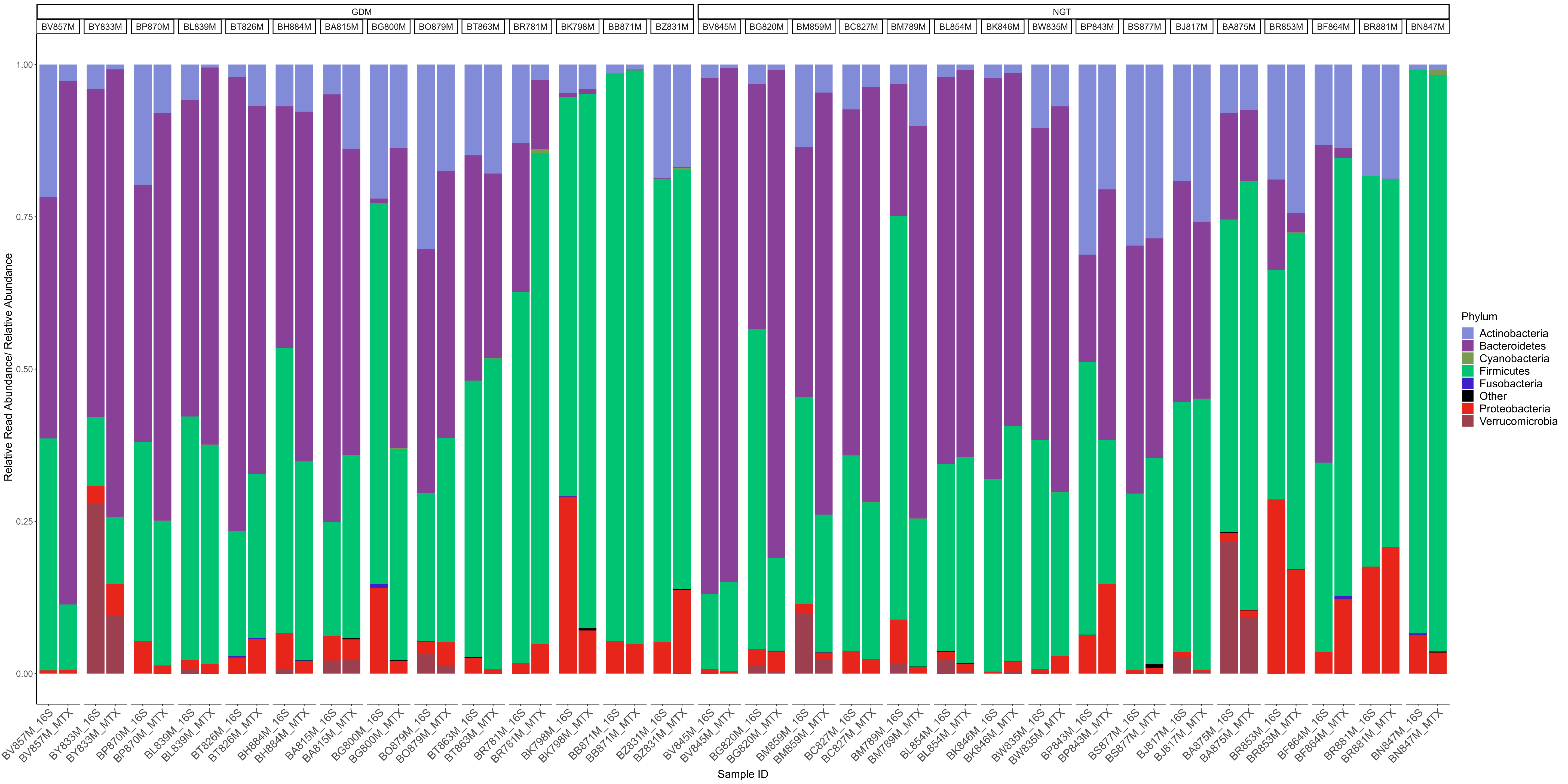

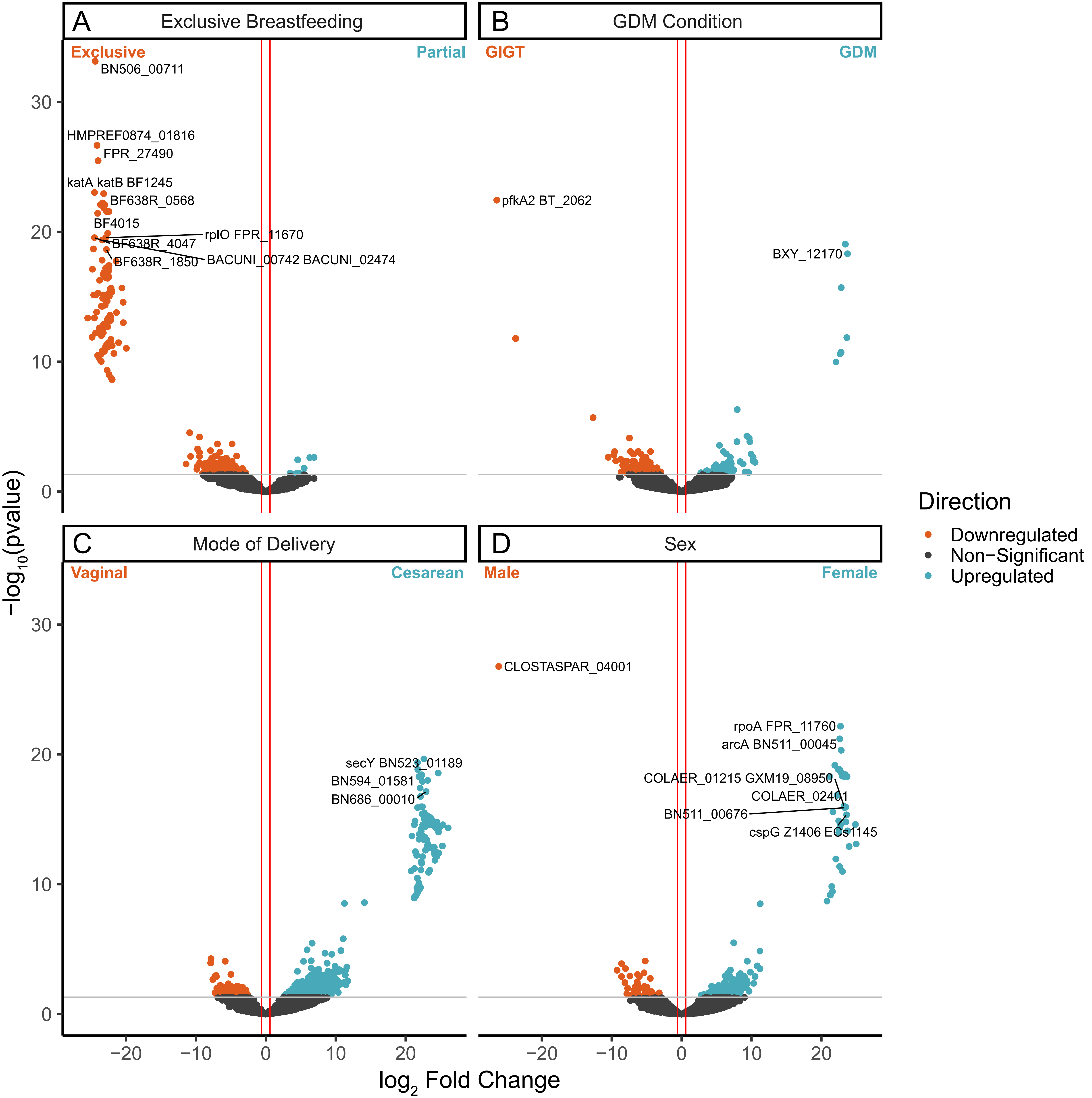

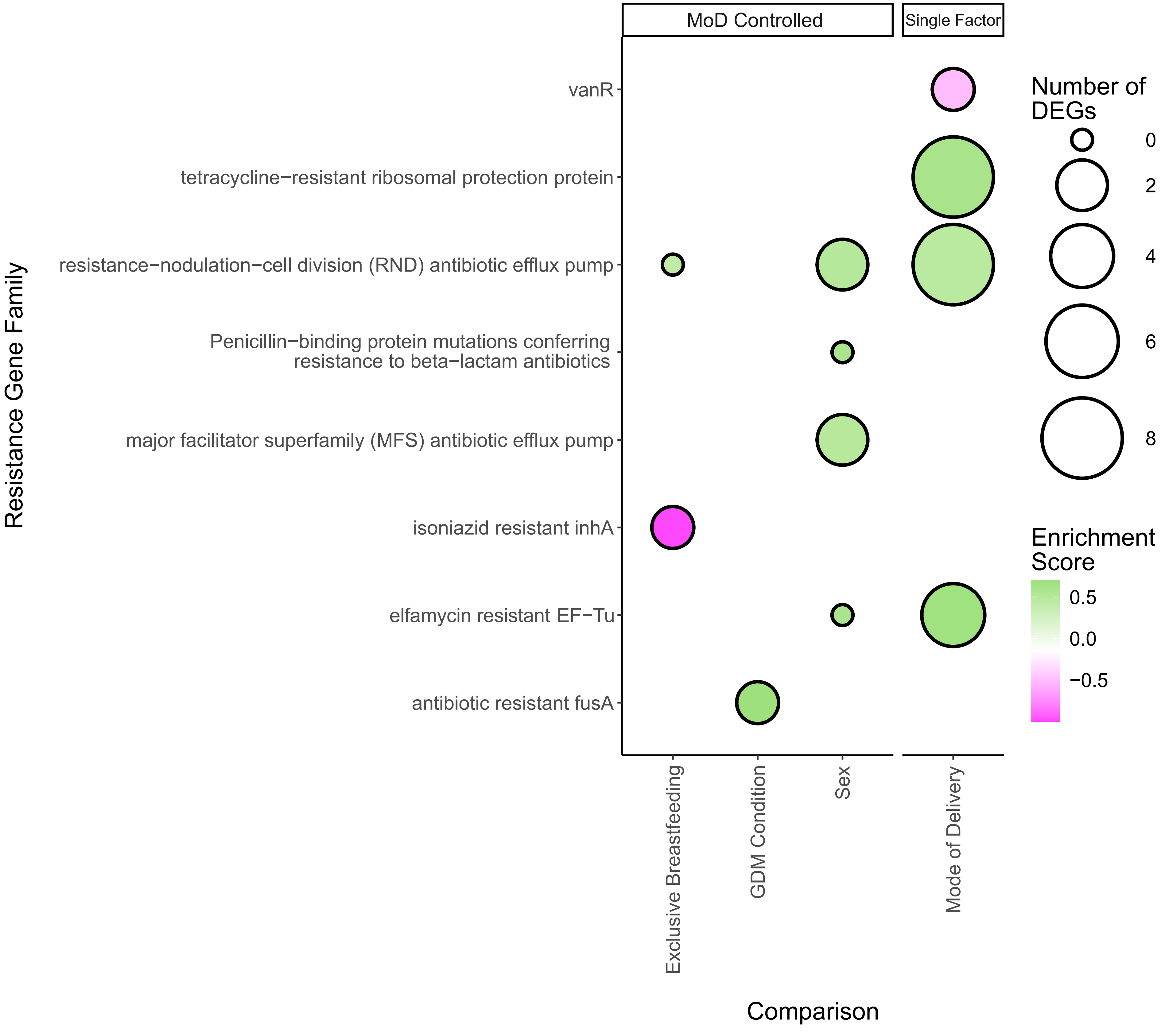
